# A protocol for rapid generation cycling (speed breeding) of hemp (*Cannabis sativa*) for research and agriculture

**DOI:** 10.1101/2022.09.12.507554

**Authors:** Susanne Schilling, Rainer Melzer, Caroline A. Dowling, Jiaqi Shi, Shaun Muldoon, Paul F. McCabe

**Author notes:** These authors contributed equally (Susanne Schilling, Rainer Melzer).

## Abstract

Hemp (*Cannabis sativa*) is a highly versatile multi-purpose crop with a multitude of applications, from textiles, biofuel and building material to high-value food products for consumer markets. Furthermore, non-hallucinogenic cannabinoids like cannabidiol (CBD), which can be extracted from female hemp flowers, are potentially valuable pharmacological compounds. In addition, hemp has high carbon sequestration potential due to its rapid growth rate. Therefore, the hemp industry is gaining more traction and breeding hemp cultivars adapted to local climate conditions or bred for specific applications is becoming increasingly important.

Here, we present a method for rapid generation cycling (speed breeding) for hemp. The speed breeding protocol makes use of the photoperiod sensitivity of *Cannabis*. It encompasses vegetative growth of the plants for two weeks under continuous light, followed by flower induction, pollination and seed development for four weeks under short-day conditions and a seed ripening phase under continuous light and water stress. With the protocol introduced here, a generation time of under nine weeks (61 days) from seed to seed can be achieved. Our method furthermore synchronises flowering time of different hemp cultivars, thus facilitating crosses between cultivars. The extremely short generation time will enable hemp researchers and breeders to perform crosses in a time-efficient way and generate new hemp cultivars with defined genetic characteristics in a shorter amount of time.

## Background

Hemp (*Cannabis sativa*) is a highly versatile crop with dozens of different applications, ranging from the fibres that can be manufactured into textiles and ropes to the highly nutritious seeds that can be processed into oil for human consumption. In addition, hemp has high carbon sequestration potential when used as hempcrete or insulation material (Ingrao et al., 2015; Schilling et al., 2021) and utilised as biofuel, hemp can act as a source of green carbon and renewable energy (Tour et al., 2010). The beneficial effects of non-hallucinogenic cannabinoids like cannabidiol (CBD) are being analysed for the treatment of various diseases, such as anxiety and depression, Parkinson’s disease, epilepsy and different types of cancers (Schluttenhofer and Yuan, 2017). Therefore, after previously being banned for decades in many countries, hemp farming and breeding are gaining more traction, with the need for cultivars adapted to the local climate or for a specific application steadily increasing.

Rapid generation cycling (speed breeding) is a technique which involves the optimisation of different environmental factors to achieve a short generation time for a given species (Ghosh et al., 2018; Watson et al., 2018). Speed breeding has recently been established for several crops including wheat (*Triticum aestivum*), canola (*Brassica napus*) and chickpeas (*Cicer arietinum*) (Ghosh et al., 2018; Watson et al., 2018). These are photoperiod-sensitive species that require long days (16 hours and above) to flower; hence they are referred to as long-day plants (Andrés and Coupland, 2012). One primary measure to achieve short generation times (rapid cycling) for long-day plants is growing them under 22 to 24 h of light per day in order to transition them from vegetative to the reproductive stage of the life cycle as quickly as possible (Ghosh et al., 2018; Watson et al., 2018).

In contrast, species that require the day length to drop below a certain critical threshold to induce flowering are termed short-day plants. Speed breeding has been established for short-day plants, like soybean (*Glycine max);* however, this requires more complex protocols, involving, for example, modification of daylength, CO2 concentration or light quality (Fang et al., 2021; Jähne et al., 2020; Nagatoshi and Fujita, 2019). Hemp was among the first plants for which day length as a critical denominator for flowering was described in the scientific literature (Tournois 1912, as cited in (Heslop-Harrison, 1957; Kobayashi and Weigel, 2007)). Meanwhile, it is well established that hemp is a short-day plant and that flowering is induced by day lengths shorter than 11 to 15 h, depending on the cultivar (Moher et al., 2021; Zhang et al., 2021). This results in long generation times: the flowering time of hemp grown in middle latitudes like in northern Europe or northern parts of North America can range from 60 to more than 100 days (Faux et al., 2013; Stack et al., 2021), and the generation time in the field is often longer than 120 days (Faux et al., 2013). This, in turn, delays breeding new hemp varieties as only one generation can be grown in the field per year.

Here, we describe a method for achieving a shortened generation time of under nine weeks (61 days) for hemp. Ten different hemp cultivars with variable flowering times under field conditions were subjected to speed breeding conditions, and for all of them, flowering time could be decreased to about 30 days. Plants cultivated with our method showed a seed set and germination rate sufficient for crosses and single seed descent lines. Our method enables rapid generation cycling for hemp, facilitating both research and breeding of this high-value crop.

## Materials and Methods

### Plant material

Hemp seeds were obtained commercially (‘Finola’, ‘Fedora17’, ‘Felina32’, ‘Santhica27’, ‘Futura75’) and from the Leibniz Institute of Plant Genetics and Crop Plant Research (IPK) seed bank in Gatersleben, Germany (KOREA (CAN22), GEORGIEN (CAN23), Kompolti-a (CAN56), Kompolti-b (CAN70), Futura (CAN69)).

### Sowing and hemp plant cultivation

Plants were grown in 1-litre pots in a soil mixture (1:1:1; John Innes No 2: Vermiculite: Perlite) to allow for soil drainage with simultaneous moisture retention. Seeds were sown directly onto the soil and placed in the dark for two days at ambient temperature (20-25 °C).

Subsequently, plants were grown for two weeks under continuous light (red-blue LEDs, HortiLED MULTI, 44W; light intensity approximately 500 μmol/s at approximately 15 cm distance to the light source) in a growth chamber at 22 to 30 °C (temperature and humidity log in Figure S1). In parallel, plants were germinated and grown under natural day length in greenhouse conditions between May and September 2019 in Dublin, Ireland.

Plants were fertilized with a commercial fertiliser (Miracle Gro, Company, NPK 25-3-14, diluted in water according to manufacturer’s instructions) once a week except for the water stress period during the ripening stage.

### Day-length cycling for optimising flower induction, seed set and ripening

After cultivation under continuous light for two weeks plants were subjected to different conditions: either short-day conditions of 12 hours light and 12 hours dark (12:12), ultra-short-day conditions of 8 hours light and 16 hours dark (8:16), long-day conditions of 16 hours light and 8 hours dark (16:8) or continuous light (24:0). Plants were kept under those conditions until they flowered and had set seeds. Thereafter, plants were exposed to continuous light for the remainder of the cultivation. After one week of continuous light, watering was suspended to put the plants under water stress to facilitate rapid seed ripening as described before for other crops (Ghosh et al., 2018).

### Flowering time, pollination and seed ripening

Plants were designated as flowering if either stigma were visible (female flowers) or single flowers were visible (male). Plants were assessed for evidence of flowering every 1-3 days. For plants cultivated in the glass house (natural light) flowering was observed two to four times a week up to day 74; afterwards, flowering was observed on days 102 and 130.

Pollen was dispersed by shaking plants bearing male flowers after anthesis or by plucking male open flowers and bringing their anthers in direct contact with female flowers. Seeds were harvested once they were hardened, and the colour had changed from green to brown. Harvested seeds were directly sown out onto soil without prior treatment.

### Silver nitrate treatment for induction of male flowers

Male flowers on female plants were induced for some plants grown under short-day conditions (12:12) using an aqueous solution of 1.5 mM silver nitrate and 6 mM sodium thiosulfate as described previously (Lubell and Brand, 2018).

The silver nitrate solution was applied using a pipette on individual inflorescence primordia (50 μl). Once plants had started the short day treatments (12 h light), silver nitrate treatment was conducted two to five times starting in 3-day intervals and suspended as soon as male flowers were visible.

## Results

### Flowering time of photoperiod-sensitive hemp cultivars can be shortened by a combination of continuous light and short-day treatments

In order to identify conditions best suited to accelerate flowering in hemp, the photoperiod-sensitive cultivars ‘Fedora17’ and ‘Felina32’ and the photoperiod-insensitive cultivar ‘Finola’ were grown under artificial light in a temperature-controlled environment for a complete life cycle from seed to seed (Figure 1, S1). Plants were germinated and initially cultivated for two weeks under continuous light to facilitate vegetative development and support the accumulation of biomass sufficient to support robust flowering. Subsequently, plants were subjected to different light treatments to simulate ultra-short days (8 hours light, 16 hours dark; 8:16), short days (12:12), long days (16:8) and continuous light (24:0) to assess how different light regimes affect flowering. Additionally, hemp plants were grown under natural light in a greenhouse to simulate a field season.

**Figure 1.**
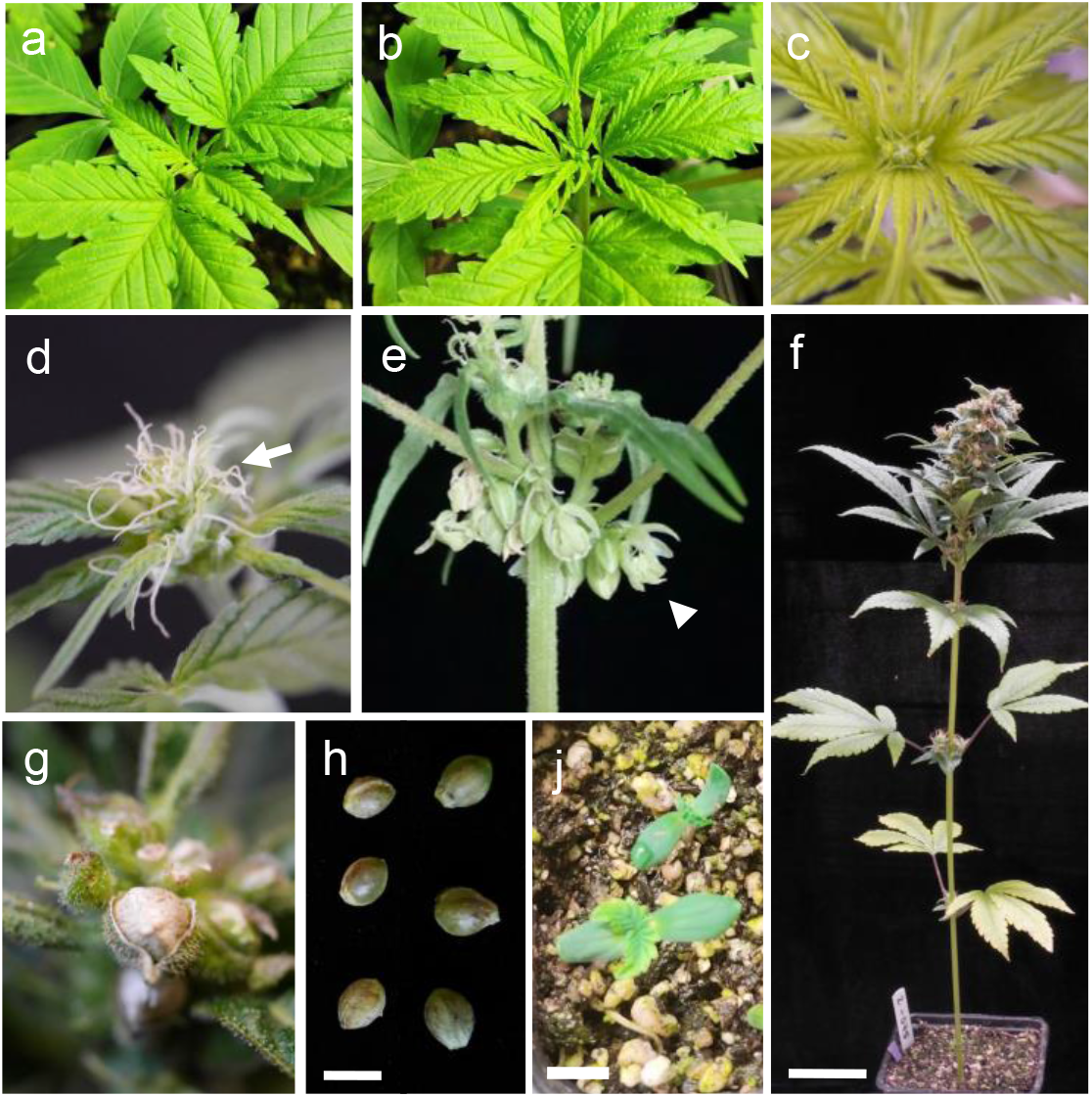
Vegetative and reproductive stages of hemp plants grown under speed breeding conditions. The sex of diecious plants cannot be morphologically determined before flowering as male (a) and female (b) plants look identical. The inflorescence of a monoecious plant becomes visible about five days before flowering (c). After flowering, female (d) and male (e) flowers are easily distinguishable, with female flowers displaying characteristic white stigmata (arrow) and male flowers containing five stamens per flower (arrowhead). A hemp plant grown under speed breeding conditions (f) shows ripening seeds after approximately 60 days (g). Ripe hemp seeds harvested 61 days after sowing (h) are able to germinate, with seedlings showing their first true leaves after a week (j). Size bars 5 cm (f) and 0.5 cm (h, j).

Alterations in flowering times were observed for different day lengths. Short days (12:12) led to a significant reduction in flowering time compared to greenhouse conditions or continuous light for both ‘Fedora 17’ and ‘Felina 32’ cultivars (Figure 2a, b). At the same time, there was no difference for the photoperiod-insensitive ‘Finola’ cultivar (Figure 2c). A further reduction of the day length to ultra-short days (8:16) did result in later flowering compared to short-day treatment for ‘Fedora17’ (Figure 2a). Photoperiod-sensitive plants grown under short-day conditions flowered approximately 44 and 70 days earlier compared to greenhouse conditions (‘Fedora17’ and ‘Felina32’, respectively, Table 1).

**Figure 2.**
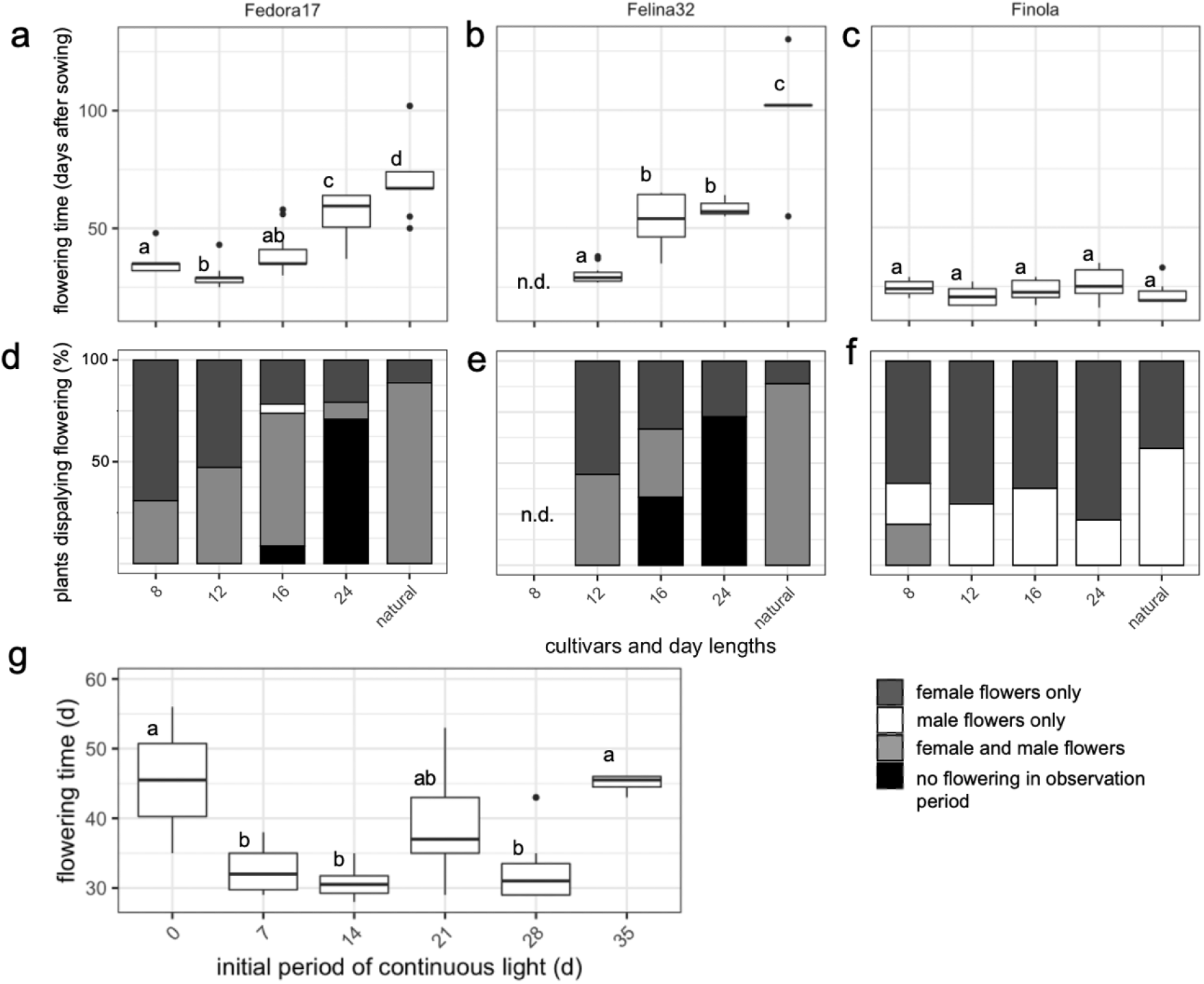
Photoperiod-sensitive hemp cultivars respond to changes in day length with alterations in flowering time and sex expression. The photoperiod-sensitive hemp cultivars ‘Fedora17’ (a) and ‘Felina32’ (b) respond with different flowering times under 8 hours light and 16 hours dark (8) (not determined for ‘Felina 32’), 12 hours light and 12 hours dark (12), 16 hours light and 8 hours dark (16) and continuous light (24). In a greenhouse with natural lighting, between May and September 2019 (natural), flowering times are highest for both cultivars. The photoperiod-insensitive cultivar ‘Finola’(c) does not change flowering time in response to different light regimes. Significance levels (p < 0.05) are indicated with letters a to d. With different day-lengths, the percentage of plants displaying flowering varied for ‘Fedora17’ (d) and ‘Felina32’ (e) but not ‘Finola’ (f). Plants showed either female-only flowers (dark grey) or male-only flowers (white) or both, male and female flowers (light grey). Percentage of plants flowering was highest at 8 hours light and 12 hours light and declining with 16 hours and continuous light, where a high percentage of photoperiod-sensitive individuals did not show any flowering at all during the observation period (black). Under ultra-short days, some ‘Finola’ plants developed both, male and female flowers. The initial vegetative growth period influences flowering time (g). Hemp plants of the cultivar ‘Fedora 17’ were grown with no initial continuous light period (0) or different ascending periods of an initial continuous period of light of 7, 14, 21, 28 and 35 days. Significance levels (p < 0.05) are indicated with letters.

**Table 1.**
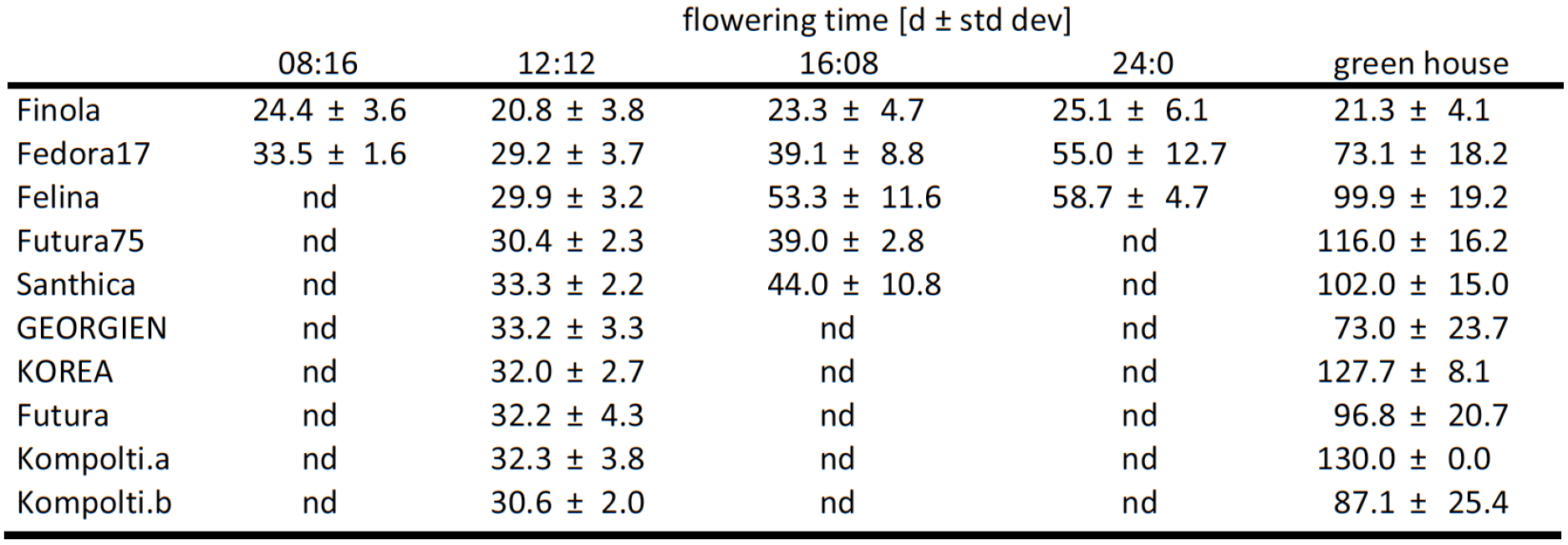
Sverage flowering time of hemp cultivars under different light regimes.

| | flowering time [d ± std dev] | | | | |
| --- | --- | --- | --- | --- | --- |
|  | 08:16 | 12:12 | 16:08 | 24:0 | green house |
| Finola | 24.4 ± 3.6 | 20.8 ± 3.8 | 23.3 ± 4.7 | 25.1 ± 6.1 | 21.3 ± 4.1 |
| Fedora17 | 33.5 ± 1.6 | 29.2 ± 3.7 | 39.1 ± 8.8 | 55.0 ± 12.7 | 73.1 ± 18.2 |
| Felina | nd | 29.9 ± 3.2 | 53.3 ± 11.6 | 58.7 ± 4.7 | 99.9 ± 19.2 |
| Futura75 | nd | 30.4 ± 2.3 | 39.0 ± 2.8 | nd | 116.0 ± 16.2 |
| Santhica | nd | 33.3 ± 2.2 | 44.0 ± 10.8 | nd | 102.0 ± 15.0 |
| GEORGIEN | nd | 33.2 ± 3.3 | nd | nd | 73.0 ± 23.7 |
| KOREA | nd | 32.0 ± 2.7 | nd | nd | 127.7 ± 8.1 |
| Futura | nd | 32.2 ± 4.3 | nd | nd | 96.8 ± 20.7 |
| Kompolti.a | nd | 32.3 ± 3.8 | nd | nd | 130.0 ± 0.0 |
| Kompolti.b | nd | 30.6 ± 2.0 | nd | nd | 87.1 ± 25.4 |

The majority of photoperiod-sensitive hemp plants grown under continuous light did not flower in the observation period (74 % and 80 % of ‘Fedora17’ and ‘Felina32’, respectively), and some individuals grown under long-day conditions (16:8) equally did not set flowers within the observation period (Figure 2d, e) while all ‘Finola’ plants flowered irrespective of day length (Figure 2f). Photoperiod-insensitive hemp plants that did flower under continuous light appeared to have a lower number of flowers compared to the plants grown under short-day conditions (data not shown). Further, biomass, height and overall stability of plants were impacted by growth conditions, with plants grown under ultra-short days being much shorter and feebler than plants grown under short days, long days and continuous light (Figure S2).

In an attempt to further shorten the flowering time, the initial period of growth under continuous light to support vegetative growth was altered. However, no significantly earlier flowering was observed if plants were moved to short-day conditions after 7 instead of 14 days (Figure 2g). Eliminating the initial vegetative growth phase resulted in significantly delayed flowering (p = 0.017). Similarly, an extension of the phase with continuous lighting to 35 days instead of 14 delayed flowering times (p < 0.01).

Subsequently, we tested whether reducing the soil volume from 1 litre to 0.2 litres would accelerate flowering time. In contrast to our expectations, smaller pots led to a significant increase in flowering time when compared to larger pots (Figure S3).

### Flowering time of hemp cultivars can be synchronised using short-day length artificial lighting

In order to verify the applicability of the developed protocol for hemp cultivars other than ‘Felina32’ and ‘Fedora17’, the flowering time of seven additional photoperiod-sensitive hemp cultivars was tested under short-day conditions and in the greenhouse. For all cultivars, flowering time under artificial lighting and short-day conditions was strongly accelerated compared to greenhouse conditions (Figure 3). The mean flowering time under short-day conditions was about 31.5 days for all photoperiod-sensitive cultivars (Table 1). Intriguingly, the flowering times of all cultivars under short-day conditions were extremely similar to each other, ranging from 29.2 to 33.3 days on average, with an average standard deviation of 3 days (Table 1, Figure 3). In contrast, flowering time under natural day lengths varied enormously between individuals and cultivars, from 50 to 130 days for photoperiod-sensitive cultivars (Table 1, Figure 3).

**Figure 3.**
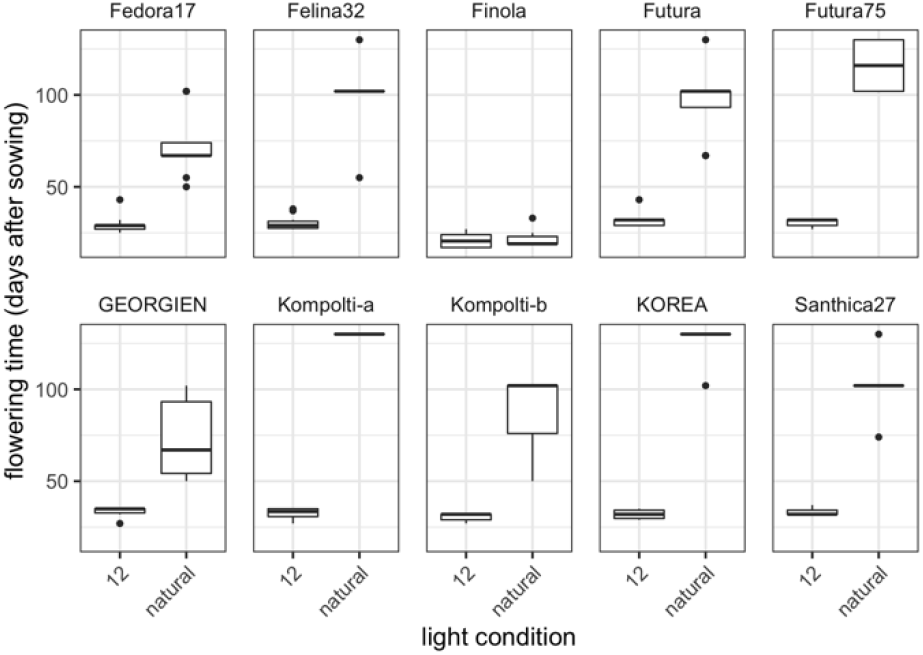
Flowering time photoperiod-sensitive hemp cultivars are lower under speed breeding conditions. Nine photoperiod-sensitive hemp cultivars, as well as the photoperiod-insensitive hemp cultivar ‘Finola’, were grown under 12 hours of light after an initial 14 days of 24 hours of light. In comparison, the same cultivars were grown in a greenhouse with natural lighting (natural). Flowering times are significantly different between treatments for all cultivars except ‘Finola’ (p < 0.01).

### Viable seeds were obtained from plants grown under speed breeding conditions

Beyond achieving a short flowering time, obtaining viable seeds is an essential part of any speed breeding protocol. After the seed set, plants were grown under continuous light and watering was stopped one week later to accelerate seed ripening. For the photoperiod-sensitive cultivar ‘Fedora 17’, the number of seeds obtained per plant varied, with the highest number of seeds observed for the plants grown under short days but no significant differences between ultra-short, short and long-day treatments (Figure 4a). ‘Fedora 17’ plants grown under continuous light did not set seeds (Figure 4a). For photoperiod-insensitive ‘Finola’ hemp plants, no significant differences in seed number were observed between day lengths, including the continuous light treatment, which overall yielded the highest number of seeds per plant (Figure 4b). All other photoperiod-sensitive cultivars tested set seeds under short-day conditions, with no significant differences between cultivars (Figure 4c).

**Figure 4.**
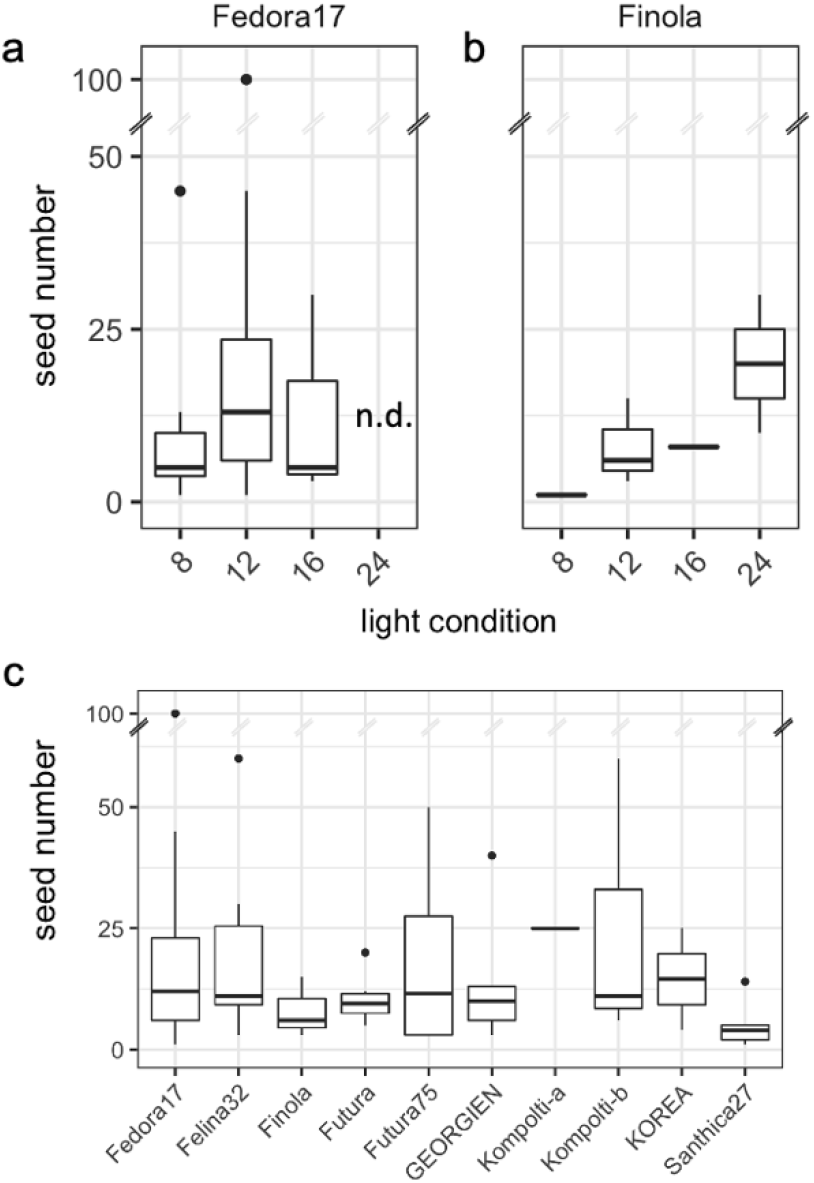
Seed yield from plants grown under speed breeding conditions. The photoperiod-sensitive cultivar ‘Fedora17’ (a) produced most seeds under short-day conditions (12), with lower yields at ultra-short (8) and long days (16). No seeds set when plants were cultivated under continuous light (24). The photoperiod-insensitive cultivar ‘Finola’ produced most seeds under continuous light (b). For both ‘Fedora 17’ and ‘Finola’ no significant differences were detected in seed numbers between conditions. Different photoperiod-sensitive hemp cultivars grown under short-day conditions produced an average of 13.2 seeds per individual with no significant differences between cultivars (c).

To assess germination rate, seeds generated from plants grown under short days conditions were placed directly on soil for germination with no prior treatment. Germination rates on average were 55 % and 86 % for ‘Fedora 17’ and ‘Felina32’ seeds, respectively (Figure S4). The fastest generation time achieved, from seed to seed, was 61 days.

### Male flowers can be induced by the application of silver nitrate

Hemp is primarily dioecious, i. e. male and female flowers develop on separate individuals. Some cultivars are monoecious, with male and female flowers developing on the same plant.

Surprisingly, we found that for the monoecious cultivars ‘Felina 32’ and ‘Fedora 17’ about 50% of the plants developed only female flowers when grown under short-day conditions, rendering these plants male sterile. In contrast, more than 80 % of the plants grown under natural light in the greenhouse developed male as well as female flowers, demonstrating that most likely this is due to the specific cultivation conditions (Figure 2d, e).

For some research or breeding applications, however, it is important to self plants. We, therefore, tested whether we can induce male flowers on female plants grown under speed breeding conditions. Silver nitrate in combination with sodium thiosulfate has been described previously for male flower induction in dioecious females (Lubell and Brand, 2018). Repeat direct application of an aqueous silver nitrate sodium thiosulfate solution to developing inflorescences of both monoecious as well as dioecious female plants resulted in the induction of male flowers approximately 14 days after the first treatment (Figure S5).

## Discussion

### A protocol to synchronise and accelerate flowering in photoperiod-sensitive hemp cultivars

Hemp (*C. sativa*) is a short-day plant and will only flower 2-4 months after sowing in field conditions in Europe and North America (Faux et al., 2013; Stack et al., 2021). Long generation times can hinder genetic research on hemp and slow down the generation of new hemp cultivars.

Here, we describe a method for rapid generation cycling (speed breeding, Figure 5) for photoperiod-sensitive hemp cultivars. The combination of an initial phase of continuous light, followed by short-day conditions (12 hours light, 12 hours darkness), provides a robust and quick transition from vegetative to reproductive development in all hemp varieties tested (Figure 1). The flowering time for photoperiod-sensitive hemp cultivars decreased dramatically under short-day conditions compared to other conditions tested (Figure 2, 3).

**Figure 5.**
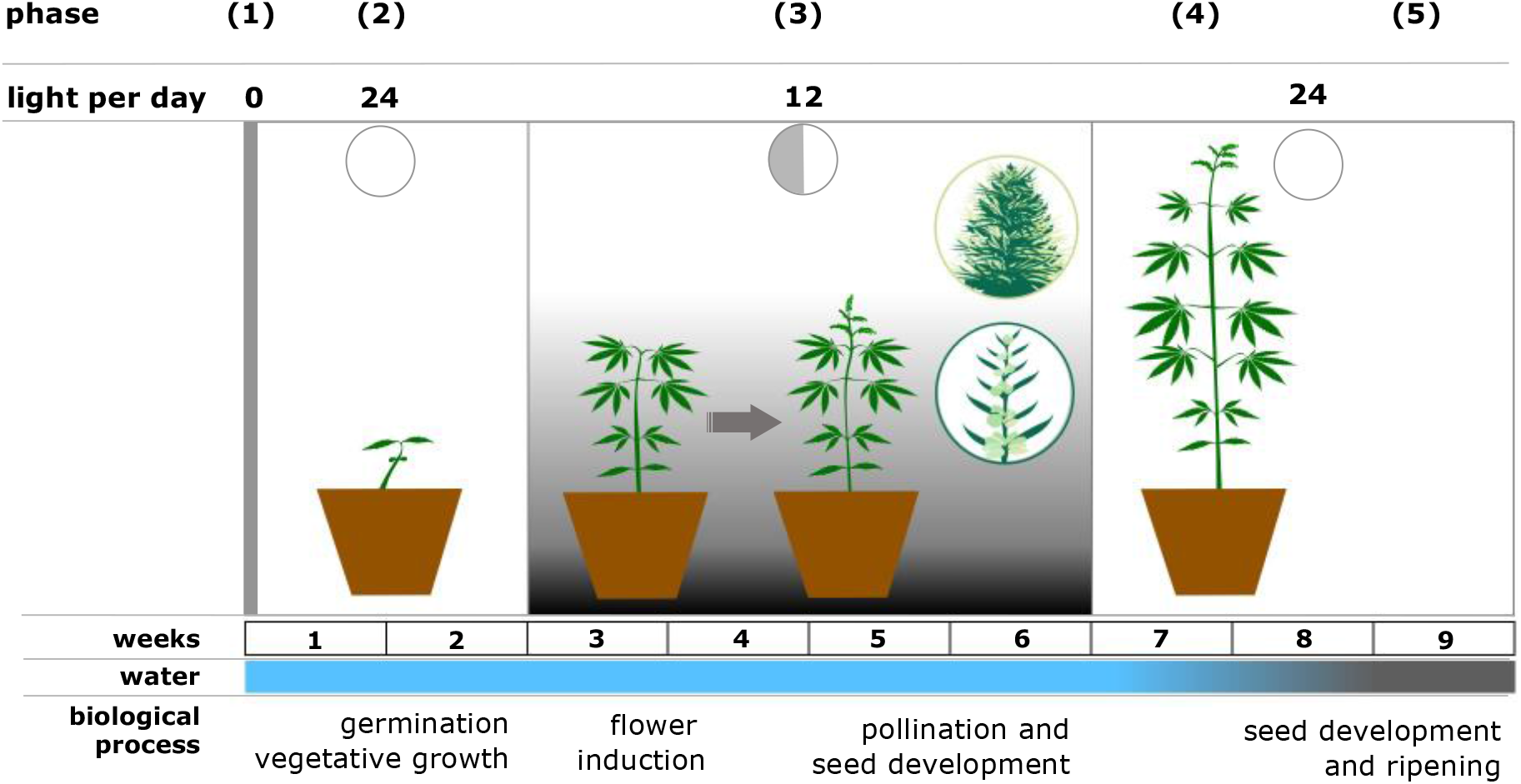
Overview of speed breeding protocol for hemp (*Cannabis sativa*). After sowing, seeds are incubated for two days in darkness (phase 1). Seedlings are then grown for two weeks under continuous light to facilitate vegetative growth (phase 2). Subsequently, hemp plants are grown under short-day conditions (12 hours light per day) for four weeks to induce flowering, pollination and seed setting (phase 3). After four weeks the plants are grown under continuous light to limit further flowering and accelerate seed ripening (phase 4). Water stress is introduced in week 8 to accelerate seed ripening (phase 5).

Hence, we propose the following protocol for rapid generation cycling for photoperiod-sensitive hemp cultivars (Figure 5):

1. Seeds are placed on soil and germinated in darkness for two days.
2. To facilitate a rapid vegetative growth of the plants, the seedlings are subjected to two weeks of continuous light.
3. To induce flowering and facilitate pollination and seed setting, plants are grown under short-day conditions (12 hours of light per day for approximately 4 weeks or until the first seeds have set).
4. To facilitate seed ripening, the plants are subjected to continuous light. Additionally, water stress is induced once seeds have reached their full size in order to accelerate seed ripening.
5. Seeds are harvested approximately 30 to 35 days after flowering when they turn brown.

With the method described here, a generation time of under nine weeks (61 days) from seed to seed can be achieved. This very short generation time can enable hemp researchers as well as breeders to perform crosses in a time-efficient way. Hence, this method can be used to create mapping populations for genetic studies and generate new hemp cultivars with defined genetic characteristics in a shorter amount of time.

### Short-day treatment synchronises the flowering time of different hemp varieties

By growing plants under short-day conditions, flowering of all hemp cultivars could be induced after a period of about 13 to 17 days under short-day conditions, even though flowering under natural light was highly variable between and even within cultivars (Figure 3, Table 1). This synchronisation enables the efficient crossing of different hemp cultivars, even if these have very distinct flowering times under natural light conditions.

It has been reported for Arabidopsis and Boechera that canalization of developmental patterns can depend on environmental conditions (Hall et al., 2007; Lee et al., 2014). It will be interesting to study what the genetic basis of this synchronisation in hemp is. Given that hemp is a short-day plant, the variation of flowering time under natural (long days) light conditions may reflect variation in the genetic background of different cultivars. Under inductive short-day conditions, the floral induction pathway may then be similar for the different cultivars, leading to similar flowering times.

### Hemp undergoes an initial vegetative period of at least 2 weeks before the competence to flower is acquired

The transition from vegetative to reproductive development in hemp can be divided into three phases: a basic vegetative phase, a photoperiod-induced phase and a flower development phase (Lisson et al., 2000a).

The basic vegetative phase is the juvenile phase, during which flowering does not occur even if day length is potentially inductive. This phase has been reported to last approximately 3 weeks or longer, depending on the cultivar (Amaducci et al., 2008; Lisson et al., 2000a). Our data agree with those observations and show that a period of continuous light during the first two weeks of growth does not delay flowering time (Figure 2g), likely because the plants have not acquired the developmental competence to flower yet. However, this initial continuous light period benefits plant growth, as it generally improves vegetative development.

The photoperiod-induced phase follows the basic vegetative phase and is day-length dependent. Under optimal light conditions, flowering initiation is assumed to be instantaneous once the plants have entered the photoperiod-induced phase (Lisson et al., 2000a). The optimal day length varies for different hemp cultivars but has been reported to be between 12 h and 14 h for most cultivars (Zhang et al., 2021). In agreement with this, flowering is readily induced at 12 h day length and delayed at 16 h. Ultra-short days do not provide an additional advantage, as flowering time does not decrease further when compared to short-day conditions (Figure 2). Under ultra-short days, the plants accumulate too little biomass and are less durable than under short days (Figure S2). It is also noteworthy that under treatment with continuous light, some plants from both photoperiod-sensitive cultivars tested flowered eventually (Figure 2), confirming earlier reports that hemp is a facultative short-day plant (Salentijn et al., 2019).

Taken together, our data and the available literature (Amaducci et al., 2008; Lisson et al., 2000a; Zhang et al., 2021) suggest that the light conditions suggested above are close to optimal for rapid induction of flowering in hemp. Seed yield is comparatively low under these conditions but should be sufficient for various breeding and research purposes such as single seed descent lines and crosses.

Interestingly, we found that short-day treatments increased the number of plants developing only female and no male flowers (Figure 2d). An effect of day length on hemp sex expression is documented (Heslop-Harrison, 1957) but has received comparatively little attention so far. Male flowers could efficiently be induced with the ethylene inhibitor silver nitrate (Figure S5). It will be interesting to study how photoperiod and hormonal pathways crosstalk to regulate sex expression in hemp.

Beyond photoperiod, growth temperature has a major effect on hemp growth, with a temperature of 26 °C to 29 °C being described as optimal (Amaducci et al., 2008; Lisson et al., 2000b). In our experiments, temperatures were around 25 °C (Figure S1). Thus, slightly higher cultivation temperatures may yield a slight further acceleration of hemp development and flowering time.

## Conclusion

We show that speed breeding in hemp is inexpensive, can be done with relatively affordable equipment, is space efficient and facilitates crossing between cultivars that diverge in flowering time under non-inductive conditions. It can serve as an efficient method to facilitate hemp breeding and research.

## Acknowledgements

SS was supported by a Postdoctoral Scholarship of the Irish Research Council and Greenlights Medicines under the Enterprise Partnership Scheme (Grant No.: EPSPD/2019/220, Enterprise partner Greenlight Medicines). CAD is supported by an Irish Research Council-Environmental Protection Agency Government of Ireland Postgraduate Scholarship (Grant No: GOIPG/2019/1987). JS is supported by a Chinese Research Council Postgraduate Scholarship (Grant No.: CSC No. 201908300031). We are grateful to James Linden and Le Roy Dowey from GreenLight Medicines for support and stimulating discussions.

## Supplementary Material

### Supplementary Figures

**Figure S1.**
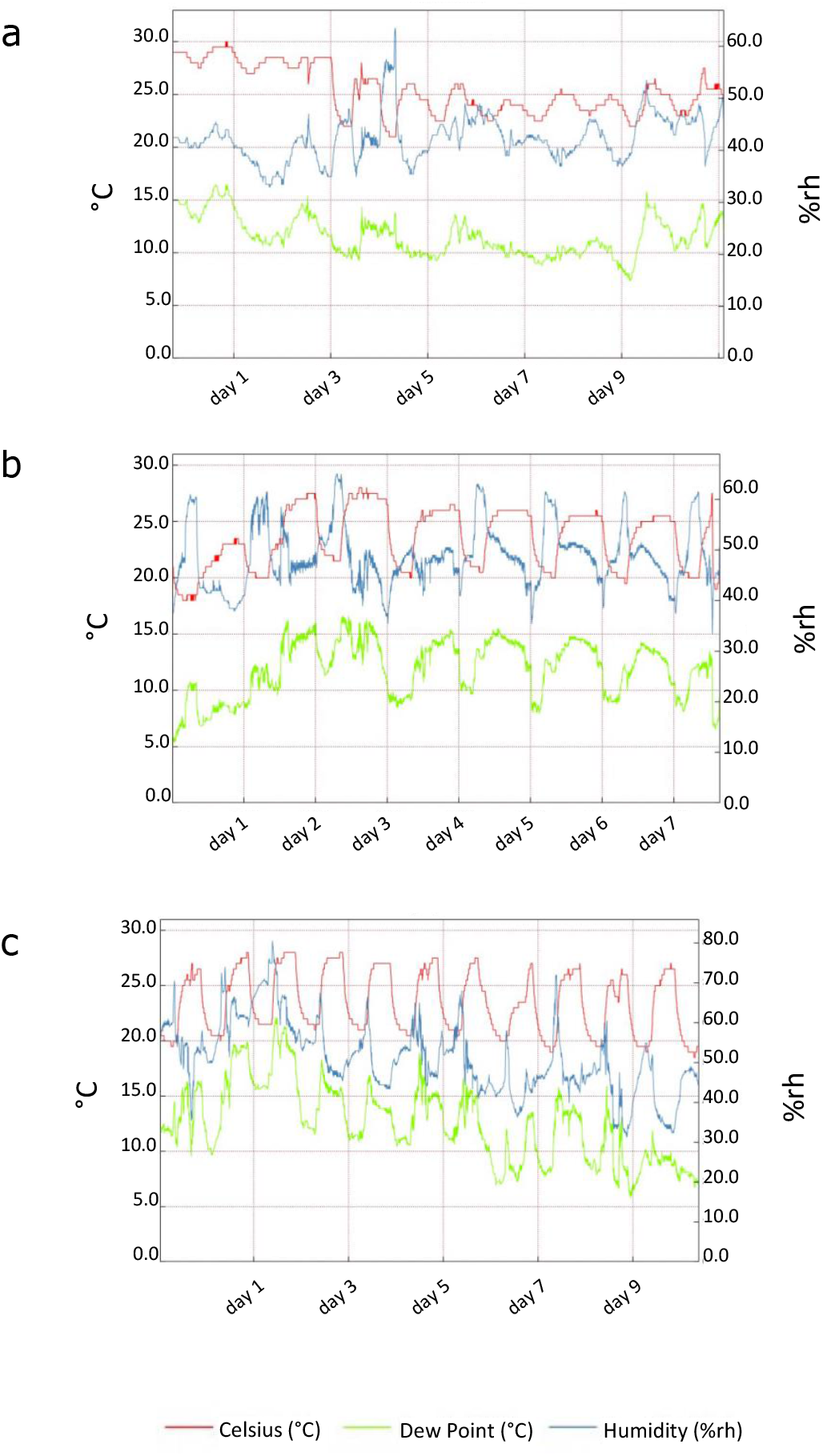
Temperature and humidity records. Measurements were recorded on 17 consecutive days (at least 5 for each light regime), which are representative of the overall experimental conditions for 24 hours of continuous light (a), 16 hours of light (b) and 12 hours of light (c).

**Figure S2.**
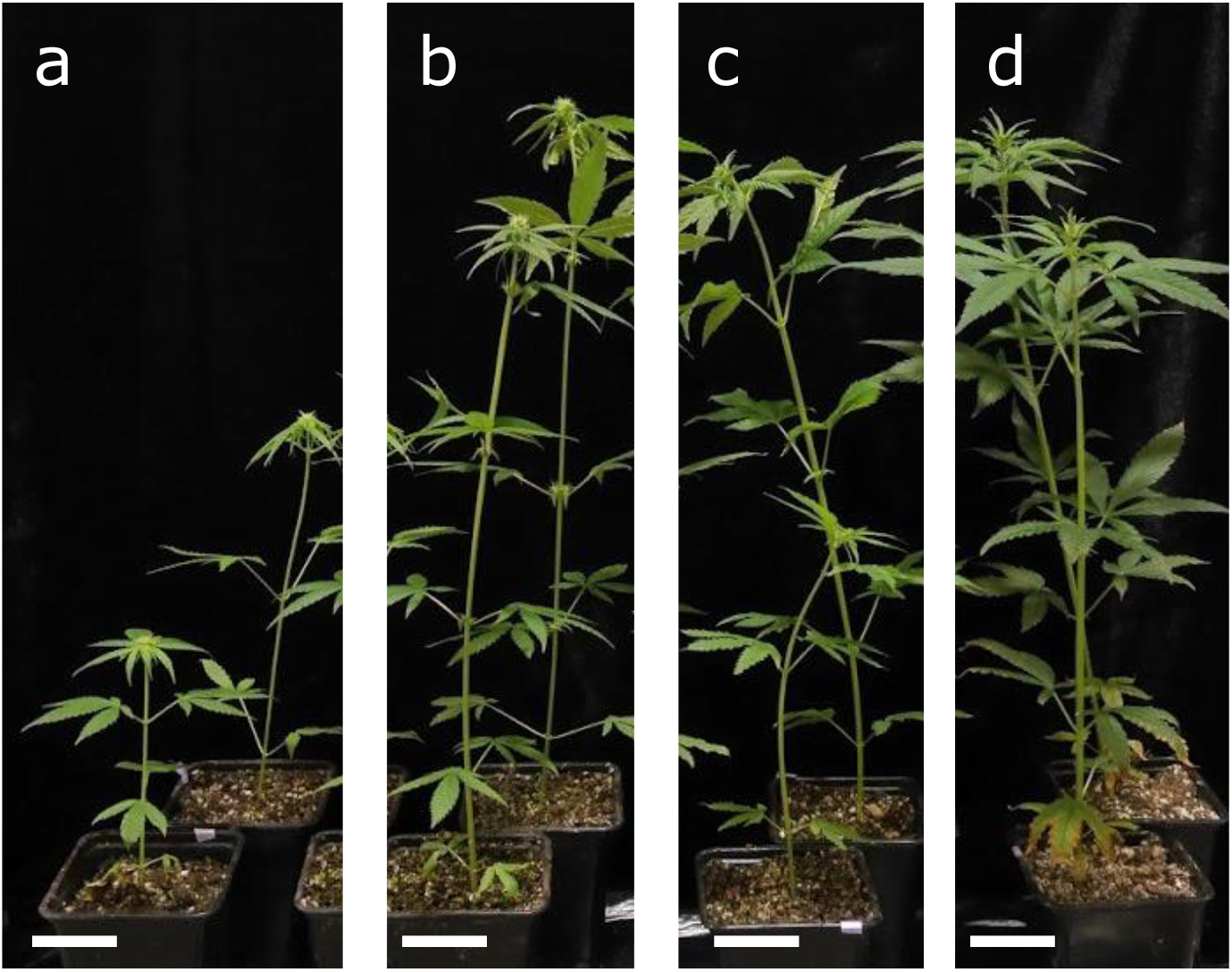
Hemp plants grown under different light conditions are different in size. Hemp plants of the cultivar ‘Fedora 17’ accumulate less biomass and are less durable if grown under ultra-short days with only 8 hours of light (a) as opposed to plants grown under 12 hours of light (b), plants grown under 16 hours of light (c) and plants grown under continuous light (d).

**Figure S3.**
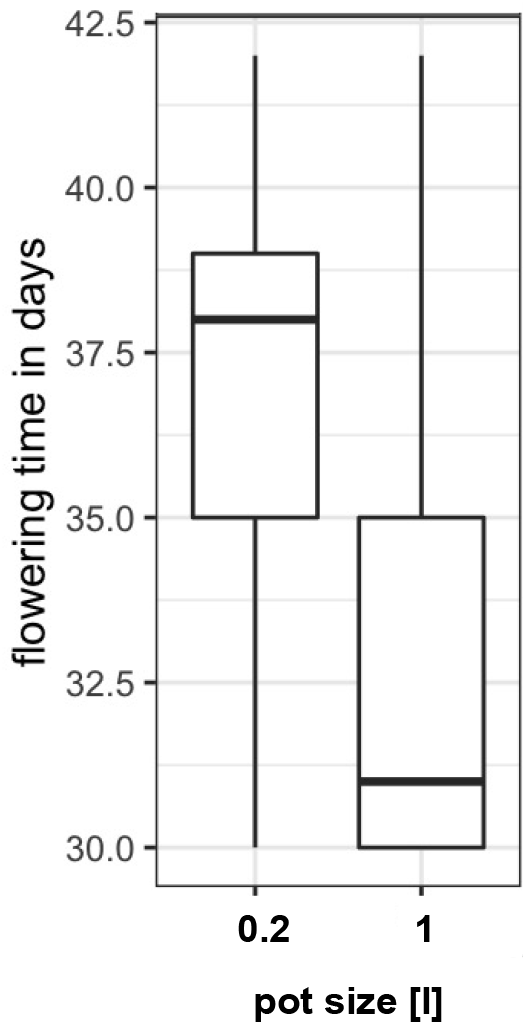
Pot size influences flowering time. Hemp plants of the cultivar ‘Fedora17’ flowered significantly later when pot size was 0.2 l as opposed to plants cultivated in 1 l pots (p = 0.00282).

**Figure S4.**
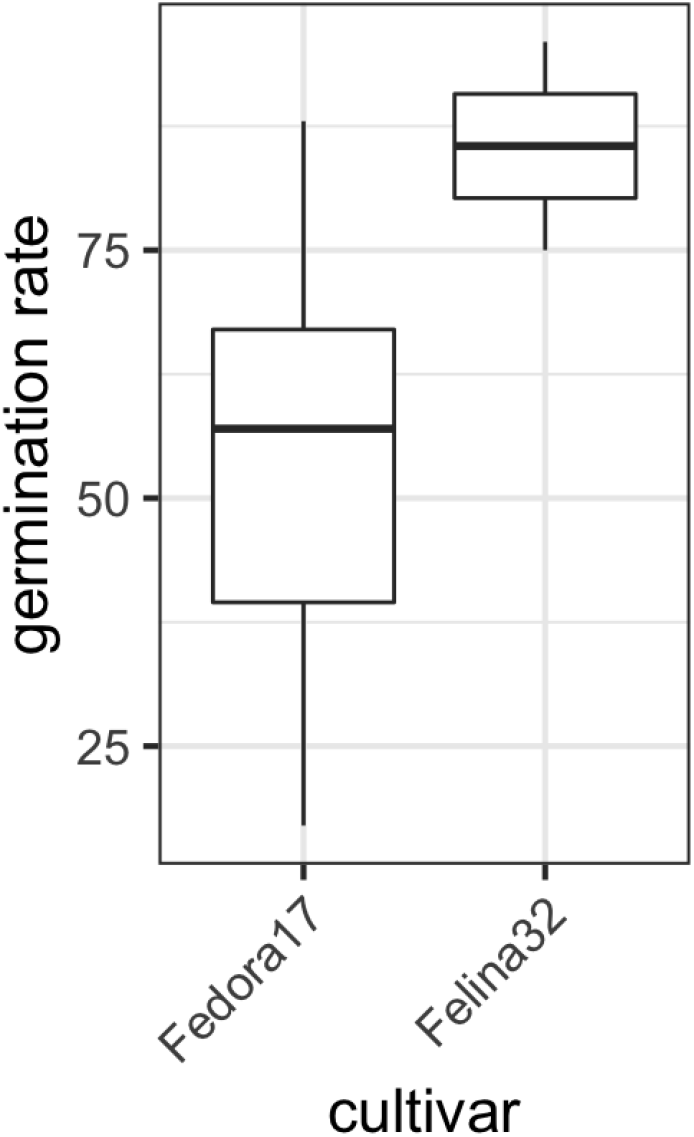
Germination rate of seeds generated with the speed breeding protocol. The germination rate of seeds was variable.

**Figure S5.**
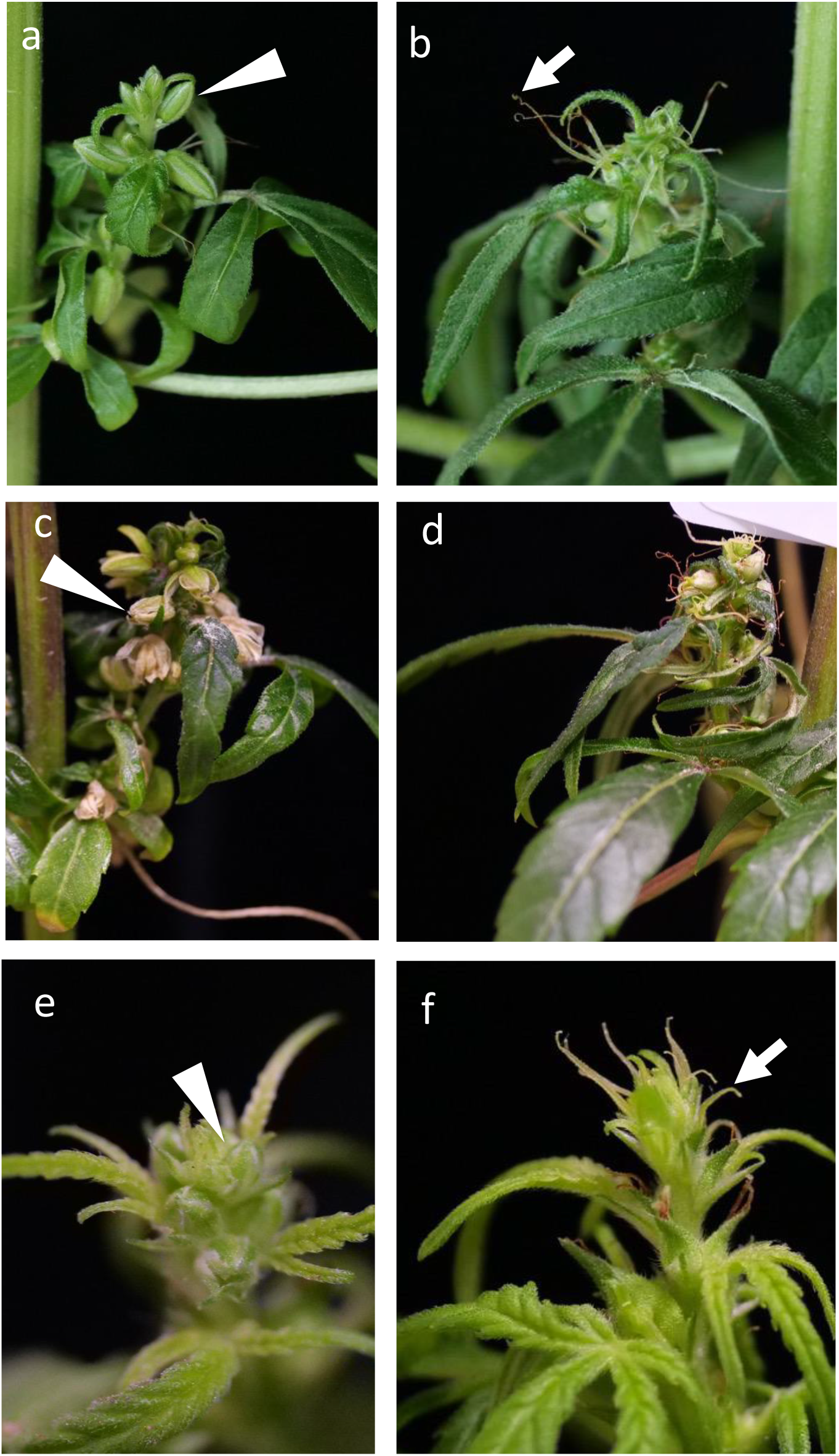
Treatment of monoecious and dioecious hemp plants with aqueous silver nitrate induces the development male flowers. Inflorescences of a monoecious (a-d) and a dioecious female (e-f) hemp plant after treatment with aqueous silver nitrate solution (a, c, e) and water control (b, d, f). Inflorescences treated with silver nitrate develop predominantly male flowers (white arrowheads). In contrast, inflorescences treated with water as a control develop predominantly female flowers (white arrow).

